# Screening for axon regeneration promoting compounds with human iPSC-derived motor neurons

**DOI:** 10.1101/2021.11.02.466937

**Authors:** Tammy Szu-Yu Ho, J. Tabitha Hees, Zhuqiu Xu, Riki Kawaguchi, Natalia P. Biscola, Daniel G Taub, Kuchuan Chen, Xirui Chen, Lee B. Barrett, Long Cheng, Christopher V. Gabel, Leif A. Havton, Daniel H. Geschwind, Clifford J. Woolf

## Abstract

CNS neurons do not regenerate after injury, leading to permanent functional deficits. Although sensory and motor neuron axons do regrow after peripheral nerve injury, functional outcome is limited due to the incomplete and slow regrowth. The lack of human-relevant assays suitable for large-scale drug screens has limited neuro-repair therapy discovery. To address this we developed a phenotypic screening strategy using human induced pluripotent stem cell-derived motor neurons to identify axon-regeneration promoting compounds and targets. The screens involve both re-plating human motor neurons on chondroitin sulfate proteoglycans and measuring regeneration responses to laser axotomy in spot cultures, and from them we identified multiple hits that promote injured axon regrowth. The top hit blebbistatin, a non-muscle myosin II inhibitor, accelerated axon regeneration and functional recovery after sciatic nerve injury *in vivo*. Human “injury in a dish” assays are suitable, therefore, to screen for therapeutic interventions that can induce or accelerate axon regeneration.

## Introduction

Neurons cannot regenerate after injury in the adult mammalian central nervous system (CNS) because of a combination of the failure to induce transcription of those genes that promote axon elongation and target innervation (Afshari et al., 2009; Giger et al., 2010; Mar et al., 2014; Renthal et al., 2020) and the presence of an extracellular and cellular environment not conducive to axonal growth (Fawcett, 2020; Filbin, 2003; Yiu and He, 2006). Although the axons of sensory and motor neurons do regenerate in the peripheral nervous system (PNS), motor functional recovery in patients with proximal nerve injury is minimal (Grinsell and Keating, 2014; Lundborg, 2000; Ma et al., 2011; Novak et al., 2011; Songcharoen et al., 2001), because of the slow pace of axonal regrowth after injury and the consequent delayed muscle re-innervation (Gordon et al., 2011; Hoke et al., 2002). CNS and PNS injuries affect millions and are devastating, yet currently, there is no approved drug to either induce regeneration of injured CNS neurons or accelerate regeneration in the PNS.

A lesion to the peripheral axons of dorsal root ganglion neurons activates intrinsic axonal growth-promoting genes (Chandran et al., 2016; Hoffman, 2010; Renthal et al., 2020), and as a consequence, a pre-conditioning peripheral nerve lesion both enhances regeneration of peripheral sensory axons (Jenq et al., 1988; McQuarrie et al., 1977) and enables regeneration of the central axons of sensory neurons in the dorsal columns beyond the lesion site in the spinal cord (Neumann and Woolf, 1999), indicating that a switch from a non-growth to an active axon growth state is a key element for both inducing regeneration in the CNS and enhancing it in the PNS. In addition, there is strong data supporting the benefit of either providing a growth-supportive and enhancing environment after nerve injury (David and Aguayo, 1981; Richardson et al., 1980) or suppressing an inhibitory one (Burnside et al., 2018; Hu et al., 2018; Schwab and Strittmatter, 2014; Tran et al., 2018; Wang et al., 2014). Although overexpressing key regeneration-driving genes can promote nerve regeneration (Fagoe et al., 2015; Ma et al., 2011; Seijffers et al., 2007), gene therapy is technically challenging and the question is whether it might be possible to promote regeneration and functional restoration after injury using small molecule drugs.

To develop drug-based therapies for promoting axon regeneration, robust human-relevant assays suitable for large scale drug screens are essential. One of the advantages of using models of human induced pluripotent stem cell (iPSC)-derived neurons is the feasibility to study both human biology and diseases and eliminate species differences. Compound screens have been performed on rodent neurons to look for molecules that enhance neurite outgrowth and promote regeneration (Al-Ali et al., 2015; Al-Ali et al., 2013; Huebner et al., 2019; Koprivica et al., 2005; Li et al., 2016; Ma et al., 2010; Usher et al., 2010), but this has not led to successful drug therapy. Modeling neurite outgrowth in human PSC-derived neurons is possible (Clarke et al., 2017; Sherman and Bang, 2018) and a screen to look for neurite growth modulating compounds in human iPSC-derived cortical-like neurons on PDL coated plates supplemented with laminin has been performed (Sherman and Bang, 2018). To mimic human neuron injury, a stretch injury model in a 96-well format using induced cortical-like neurons was developed (Sherman et al., 2016). However, there is currently no axotomy assay in human iPSC-derived neurons suitable for high-throughput screening.

Our goal here was both to develop a screening funnel that could identify targets that promote axon regeneration, by detecting the actions of unbiased bioactive target-annotated small molecules, and that has the high throughput characteristics necessary for future screens against large, chemically diverse libraries. Here we assessed the effect of bioactive small molecule target-annotated compounds on neurite outgrowth as a means of identifying new targets regulating regeneration, as well as compounds with potential for repurposing or drug development. Our strategy was to develop several different human iPSC-derived motor neuron regrowth phenotypic assays suitable for high-content screening. First, we screened for compounds that promote neurite outgrowth of re-plated motor neurons grown on chondroitin sulfate proteoglycans, a substrate selected to mimic an inhibitory extracellular matrix environment. For secondary screens, we both used laser injury of established axons in a human neuron spot culture assay and measured the dynamic outgrowth over days of live replated neurons after drug treatment. The final step of the screening funnel was to test the top hit *in vivo* in a mouse injury model.

From a screen of 4557 bioactive compounds we identified multiple novel hits whose action in promoting regeneration are reported here for the first time. We also found some hits consistent with previous studies in rodent models. The hit with greatest neurite outgrowth promotion in freshly dissociated neurons and regrowth of established axons after injury was blebbistatin, a non-muscle myosin II inhibitor that has been found before to promote neurite outgrowth in *in vitro* rodent primary neurons (Hur et al., 2011). Blebbistatin increased the neurite outgrowth of human induced motor neurons on CSPG more than 10-fold. This effect is much greater than known growth promotors, like ROCK inhibitors (Chan et al., 2005; Fournier et al., 2003; Hiraga et al., 2006; Lingor et al., 2008). Furthermore, the growth promoting effects of blebbistatin in the human *in vitro* injury model predicted axon regeneration in a mouse *in vivo* injury model.

## Results

### Phenotypic screen with human motor neurons to identify pro-regenerative compounds

We designed a screening funnel to identify target-annotated compounds that promote axon regeneration using human-induced pluripotent stem cell (iPSC)-derived motor neurons (Figure 1A). The primary screen, conducted with a bioactive compound library set, utilized a high-content neurite outgrowth promotion phenotypic assay. Dose response confirmation was performed on hits from the primary screen, followed by secondary screens utilizing an axonal injury and dynamic growth assay, followed by target validation by knockdown. Finally, we conducted *in vivo* studies using a mouse sciatic nerve injury model on the most promising hit to establish if the *in vitro* screen is predictive in revealing pro-regenerative targets for nerve repair.

**Figure 1.**
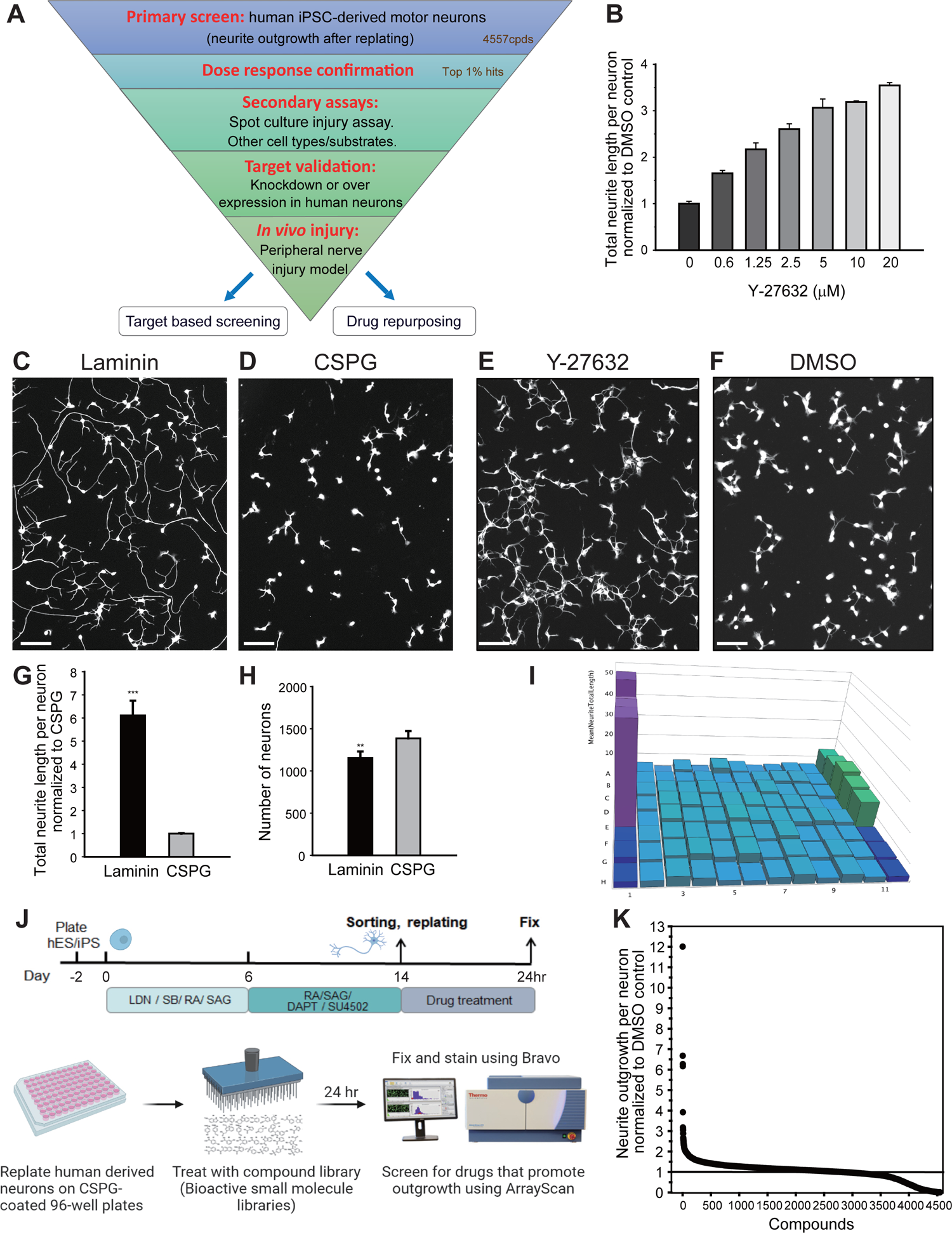
High-throughput phenotypic screen in human iPSC-derived motor neurons to identify pro-regenerative compounds. (A) Screening funnel illustrating the screening strategy. (B) Dose response curve of Y-27632, a positive control for the primary screen on neurite outgrowth (total neurite length per neuron). (C-D) Human iPSC-derived motor neurons cultured on laminin, a positive control substrate (C), or CSPG (D) for 24 hours after re-plating. Neurons stained with an antibody against βIII-tubulin. Scale bar, 50μm. (E-F) Human motor neurons treated with Y-27632 (positive control, E) or DMSO (negative control, F) for 24 hours after re-plating on the CSPG substrate. Neurons stained with anti-βIII-tubulin antibody. Scale bar, 50μm. (G) Effects of laminin vs CSPG on neurite outgrowth 24 hours after re-plating. (***P=0.000056, unpaired two-tailed t-test, n=5 for each group). (H) Number of neurons on laminin vs CSPG fixed, stained and quantified 24 hours after re-plating. (**P=0.002, unpaired two-tailed t-test, n=5 for each group). (I) A 3D histogram of a 96-well screening plate with positive (purple, laminin; green, Y-27632) and negative (dark blue, DMSO) controls. Test compounds (light blue) are in columns 2-11. Z axis shows total neurite length per neuron after compound treatment for 24 hours. (J) Primary screening strategy: Motor neurons were differentiated from human iPSC as illustrated. After differentiation, the neurons were re-plated on CSPG coated 96-well plates, treated with compounds for 24 hours, fixed, stained and imaged using the ArrayScan Screening System. (K) Primary screen results from 4557 compounds. Each dot represents a molecule. Data represented as mean ± SD.

The primary screen comprised determining axon growth 24 hours after re-plating motor neurons in wells on a substrate composed of chondroitin sulfate proteoglycans (CSPG) – a major component of the extracellular matrix in the central nervous system after injury (Yiu and He, 2006), to mimic regrowth in an injured environment. A positive control for the primary screen was laminin, a permissive substrate that promoted an increase in total neurite length per neuron of 6-fold on average (p<0.001) when compared with growth on CSPG (Figure 1C, D and G). A small molecule positive control for the screen was Y-27632 a *Rho*-associated, coiled-coil-containing protein kinase (ROCK) inhibitor that promotes axon regeneration in rodent models (Chan et al., 2005; Fournier et al., 2003; Lingor et al., 2008). In pilot experiments we found that Y-27632 promotes outgrowth of re-plated human motor neurons (Figure 1B, E and F) with a 3.5-fold increase in average total neurite length per neuron at 20μM when compared with a DMSO control, p<0.001. Figure 1B shows the dose response curve for Y-27632. A 3D histogram of a 96-well screening plate from the primary screen with the positive (purple laminin and green Y27632) and negative controls (dark blue, CSPG and DMSO) is shown in Figure 1I. The middle columns (light blue) are growth responses to a set of small molecules from a bioactive library.

The overall strategy for this primary screen is shown in Figure 1J. Human motor neurons were differentiated from iPSCs utilizing a protocol with a series of small molecules: dual SMAD inhibition to neuralize the iPSCs, FGF and Notch signaling inhibition to accelerate neuronal differentiation, and activation of retinoic acid and Sonic Hedgehog signaling to pattern and specify motor neuronal fate (Klim et al., 2019). After the differentiation, the motor neurons underwent an axonal injury surrogate by re-plating them from the differentiation plate into a 96-well plate on CSPG. After treating neurons with compounds from the library for 24 hours, we identified those that promoted neurite outgrowth, using βIII-tubulin and Hoechst staining, with imaging and quantification with an Arrayscan (Figure 1J). To quantify neurite outgrowth induced after the re-plating, the average neurite length per neuron after compound treatment was normalized to the DMSO control.

We screened several bioactive small molecule libraries composed of target-annotated compounds (see Methods) to both identify pathways regulating regeneration and compounds with potential for drug development/repurposing. A total of 4557 compounds were screened and we observed hits that promoted neurite outgrowth (fold increase in neurite length>1) as well as hits that inhibited outgrowth (fold increase in neurite length<1). The top 1% of the growth positive hits had more than a 2-fold increase in neurite outgrowth when compared with the negative control, 0.15% DMSO. The primary screen results from the replated human motor neurons on CSPG are shown in a scatter plot, where each dot represents a small molecule (Figure 1K).

### Primary screen results reveal pro-regeneration target pathways

Table 1 lists the top 20 hits from the primary screen, of which blebbistatin, a non-muscle myosin II inhibitor, had the greatest axon growth promoting action. As shown in Figure 1K, blebbistatin increased the outgrowth of human motor neurons after re-plating 12-fold when compared with the DMSO control, which is substantially greater than both our positive controls (laminin and Y-27632 a ROCK inhibitor) and indeed all other hits. Figure 2A and B show images from a screening plate of neurons treated with blebbistatin that extend more neurites than the DMSO control, and Figure 2C shows the dose response curve of blebbistatin in the assay (red line indicates effect of Y-27632 at 20μM).

**Figure 2.**
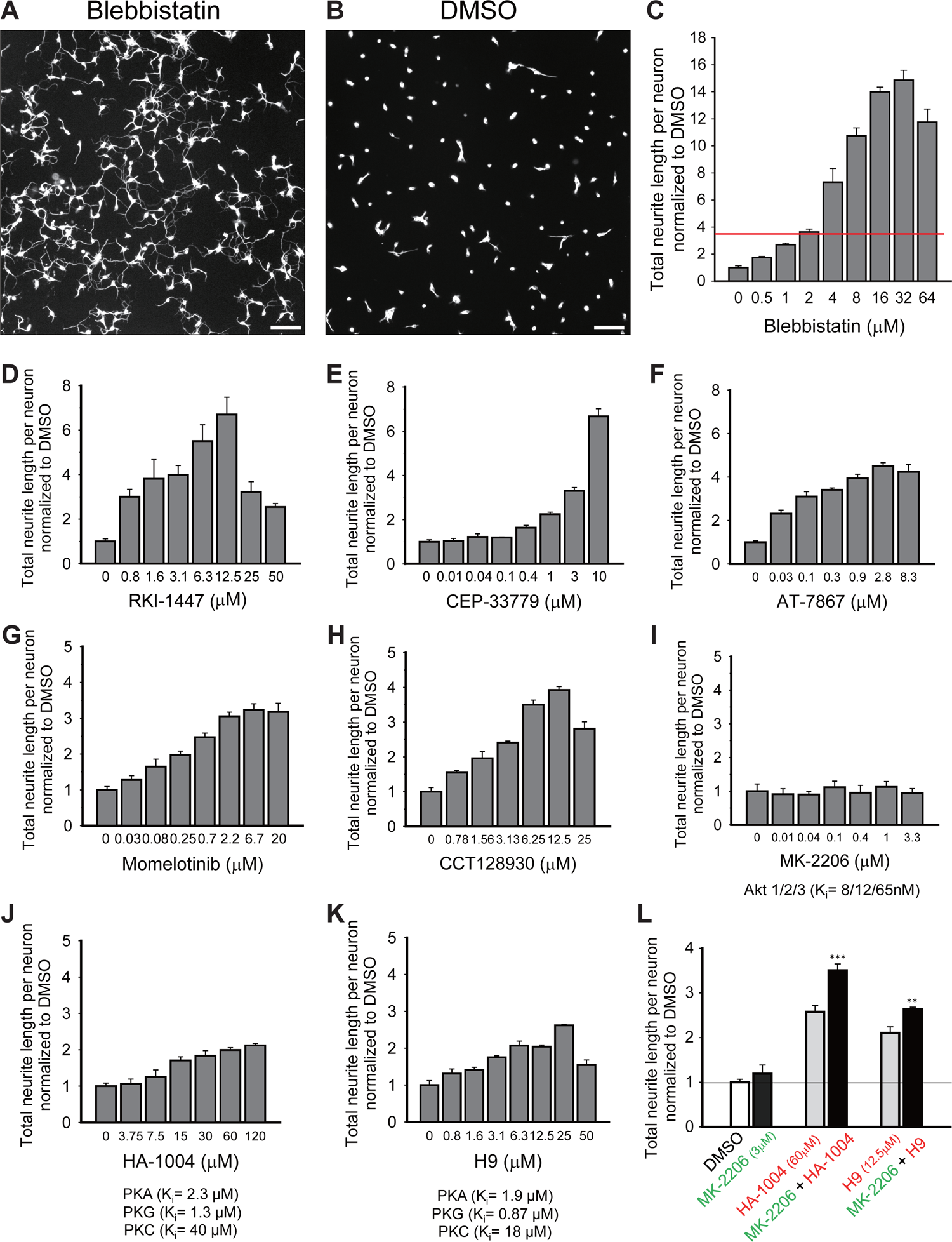
Dose response curves of top hits from primary screen. (A-B) Human motor neurons treated with 25μM blebbistatin (A) or 0.15% DMSO negative control (B) for 24 hours after re-plating on CSPG. Scale bar, 100μm. (C-H) Dose response curves of top 6 hits: blebbistatin (C), RKI-1447 (D), CEP-33779 (E), AT-7867 (F), Momelotinib (G), and CCT128930 (H), showing neurite outgrowth promotion after 24 hour treatment. (I-K) Dose response curves of an Akt inhibitor, MK-2206 (I) and PKA inhibitors, HA-1004 (J) and H9 (K). (L) Combinatorial treatment of Akt and PKA inhibitors shows better outgrowth enhancement than treatment with single inhibitors. (***P=0.0001, **P=0.0028, unpaired two-tailed t-test, n=4 for each group). Data represented as mean ± SD.

**Table 1:**
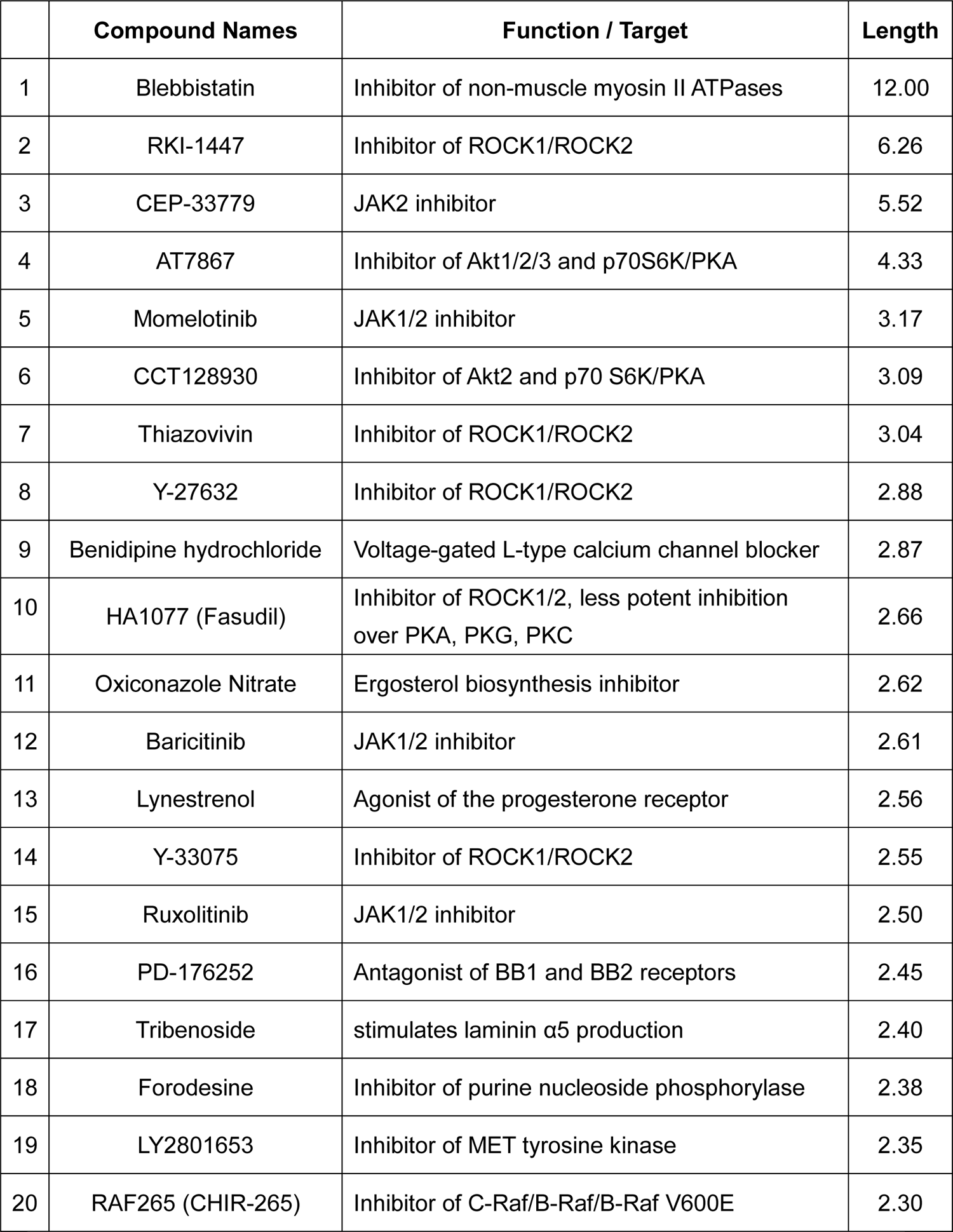
Top 20 hits from the primary screen.

|  | Compound Names | Function / Target | Length |
| --- | --- | --- | --- |
| 1 | Blebbistatin | Inhibitor of non-muscle myosin II ATPases | 12.00 |
| 2 | RKI-1447 | Inhibitor of ROCK1/ROCK2 | 6.26 |
| 3 | CEP-33779 | JAK2 inhibitor | 5.52 |
| 4 | AT7867 | Inhibitor of Akt1/2/3 and p70S6K/PKA | 4.33 |
| 5 | Momelotinib | JAK1/2 inhibitor | 3.17 |
| 6 | CCT128930 | Inhibitor of Akt2 and p70 S6K/PKA | 3.09 |
| 7 | Thiazovivin | Inhibitor of ROCK1/ROCK2 | 3.04 |
| 8 | Y-27632 | Inhibitor of ROCK1/ROCK2 | 2.88 |
| 9 | Benidipine hydrochloride | Voltage-gated L-type calcium channel blocker | 2.87 |
| 10 | HA1077 (Fasudil) | Inhibitor of ROCK1/2, less potent inhibition over PKA, PKG, PKC | 2.66 |
| 11 | Oxiconazole Nitrate | Ergosterol biosynthesis inhibitor | 2.62 |
| 12 | Baricitinib | JAK1/2 inhibitor | 2.61 |
| 13 | Lynestrenol | Agonist of the progesterone receptor | 2.56 |
| 14 | Y-33075 | Inhibitor of ROCK1/ROCK2 | 2.55 |
| 15 | Ruxolitinib | JAK1/2 inhibitor | 2.50 |
| 16 | PD-176252 | Antagonist of BB1 and BB2 receptors | 2.45 |
| 17 | Tribenoside | stimulates laminin $\alpha 5$ production | 2.40 |
| 18 | Forodesine | Inhibitor of purine nucleoside phosphorylase | 2.38 |
| 19 | LY2801653 | Inhibitor of MET tyrosine kinase | 2.35 |
| 20 | RAF265 (CHIR-265) | Inhibitor of C-Raf/B-Raf/B-Raf V600E | 2.30 |

Consistent with these findings, blebbistatin promotes neurite outgrowth in avian and rodent primary neurons and in human derived cortical-like interneurons (Hur et al., 2011; Rosner et al., 2007; Sherman and Bang, 2018). However, the drug has not been identified before in human neuron injury models.

The top 20 hits include several with an action on previously reported pre-regeneration pathways, indicating that the primary screen replicates other axon growth assays. This includes ROCK inhibitors, with five ROCK inhibitors amongst the top 20 hits, one of which was the positive control Y-27632. That ROCK inhibitors increase axon growth has been extensively studied (Chan et al., 2005; Fournier et al., 2003; Hiraga et al., 2006; Lingor et al., 2008). Among the ROCK inhibitors, RKI-1447 had the most potent effect on outgrowth promotion (Table 1 and Figure 2D). Recent reports reveal that RhoA, the immediate upstream target of ROCK, regulates cytoskeleton through non-muscle myosin II, and that the physical target of RhoA in axon growth is non-muscle myosin II (Dupraz et al., 2019). This suggests that ROCK inhibitors may promote axon regeneration by inhibiting non-muscle myosin II. Inhibiting non-muscle myosin II directly, as blebbistatin does, could though, be more advantageous than inhibiting ROCK, which will also impact other effectors downstream of ROCK. Notably, we observed a much greater promotion of neurite outgrowth after treatment with blebbistatin, the non-muscle myosin II inhibitor, than with any of the five ROCK inhibitor hits.

Another group of top hits in the primary screen were voltage-gated calcium channel blockers, especially those with a vasodilator action. These include benidipine hydrochloride (dihydropyridine), an L-, T-, and N-type calcium channel blocker used for treating hypertension (Table 1), and nicardipine hydrochloride, also an L-type calcium channel blocker and antihypertensive drug, which promoted motor neuron neurite outgrowth 2.26-fold. We earlier found in primary rodent neurons, that the L-type calcium channel blocker diltiazem (benzothiazepine) promotes axon regeneration both in rat embryonic cortical neurons and adult primary mouse dorsal root ganglion neurons, as well as in induced human sensory neurons (Huebner et al., 2019).

Four JAK inhibitors were among the top 20 hits (Table 1) and the dose response curves of these are shown in Figure 2E, G and Supplementary Figure 1A and B. That four drugs from an unbiased compound library screen that act on the same target pathway were top hits, make JAK inhibition a compelling neurite growth promotion pathway. AT7867 and CCT128930, both inhibitors of Akt1/2/3 and p70S6K/PKA, represent novel hits/targets for neurite outgrowth (Table 1; dose response curves of AT7867 and CCT128930 are shown in Figure 2F and H). To test which of the two pathways are responsible for the outgrowth promotion, we studied the individual pathways with more specific inhibitors. When motor neurons were treated with specific Akt inhibitors, no increase in neurite outgrowth was detected (Figure 2I). Although PKA inhibition did promote neurite outgrowth to some extent (Figure 2J and K), the effects were less potent than the combined inhibition of both Akt1/2/3 and PKA (Figure 2L). These data suggest that inhibition of both Akt1/2/3 and p70S6K/PKA is required for promoting human motor neuron axonal outgrowth.

### Dynamic effects of pro-regenerative hits in human motor neurons

In the large-scale primary screen we treated human motor neurons with compounds for a fixed 24 hour period and recorded neurite growth at this time. To study dynamic drug effects of hits and for a longer duration, we developed a secondary assay where we tracked the responses of live motor neurons to the drug treatment using an IncuCyte S3 (Essen BioScience), which comprises an automated microscope inside an incubator, and this allowed us to trace neurites without a reporter or label and image and analyze neuron cultures over time (five days) such that we could measure real-time responses to drug treatment. Blebbistatin dramatically promoted neurite outgrowth over the 5 days of treatment when compared with DMSO control (Figure 3A, B and Supplementary movies 1 and 2). Figure 3C shows the effects of the top 10 hits from our primary screen on dynamic human motor neuron outgrowth after re-plating, at working doses established from the dose response curves.

**Figure 3.**
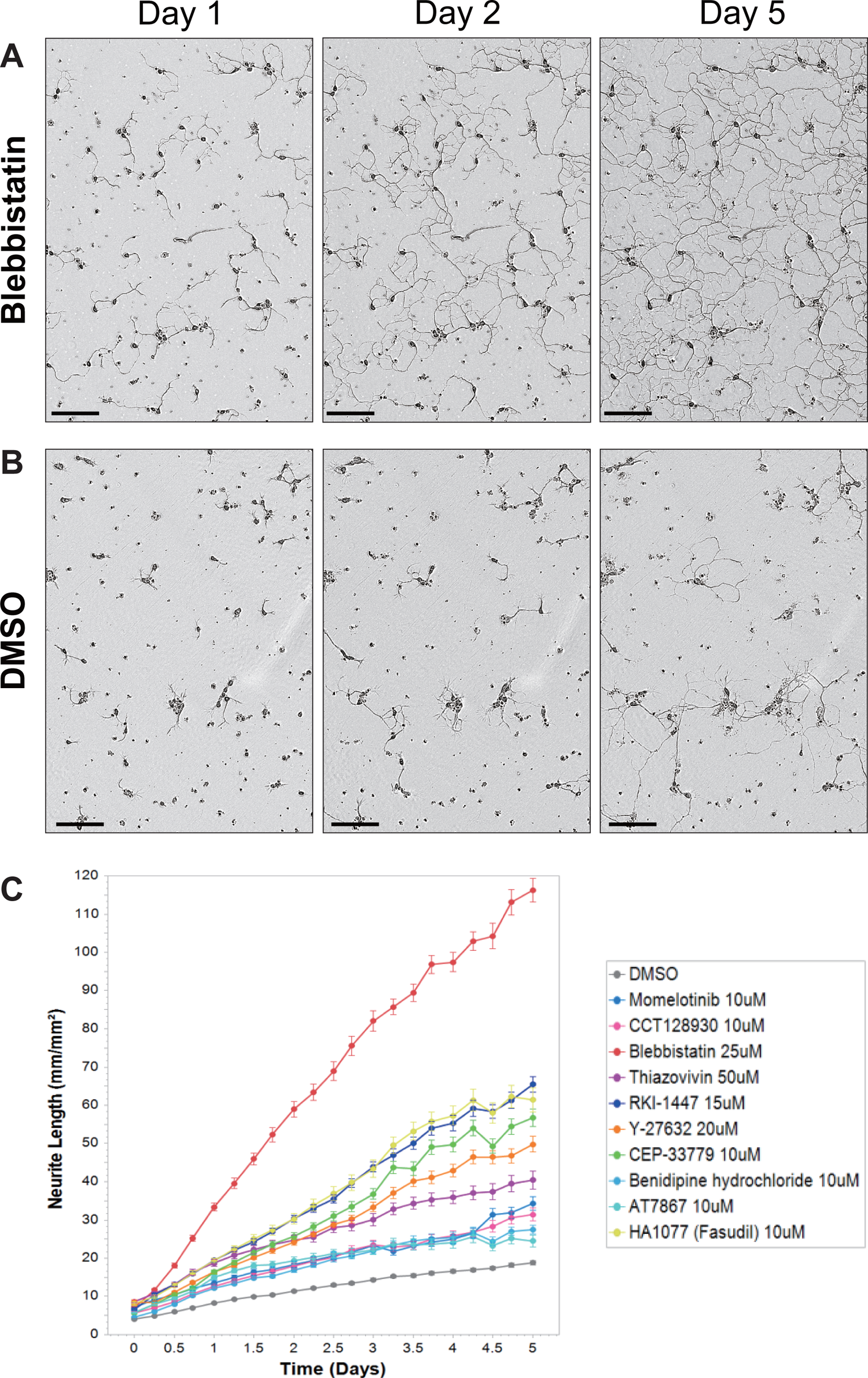
Dynamic changes in human motor neurons produced by hits. (A-B) Images of live human motor neuron taken at Day 1, Day 2 and Day 5 after 25μM blebbistatin treatment (A) or DMSO treatment (B). Scale bar, 100μm. (C) Quantification of neurite length after drug treatment over time. The top 10 hits from the primary screen were used in the dynamic re-plating assay, neurons were imaged and analyzed every 6 hours over 5 days. Data represented as mean ± SEM.

### A human neuron *in vitro* injury assay to study axon regrowth

The primary and secondary dynamic growth screens used the re-plating of differentiated neurons from a culture well as an injury surrogate. While this was efficient, reliable and robust, and therefore suitable for high-throughput primary screening, it is a harsh condition with the severing of all neuronal processes. To be closer to a patient injury situation, we developed a human neuron spot culture laser cutting assay (Figure 4A). We cultured a high density of human induced motor neurons as a dot or spot (120,000-150,000 cells per 1-7 μl/spot). Axons grow radially outward from the center of the spot (cell bodies) (Figure 4G). After growing for 7 days, a defined set of long axons projecting in the same direction could then be severed by an infrared laser (Stiletto, Hamilton Thorne).

**Figure 4.**
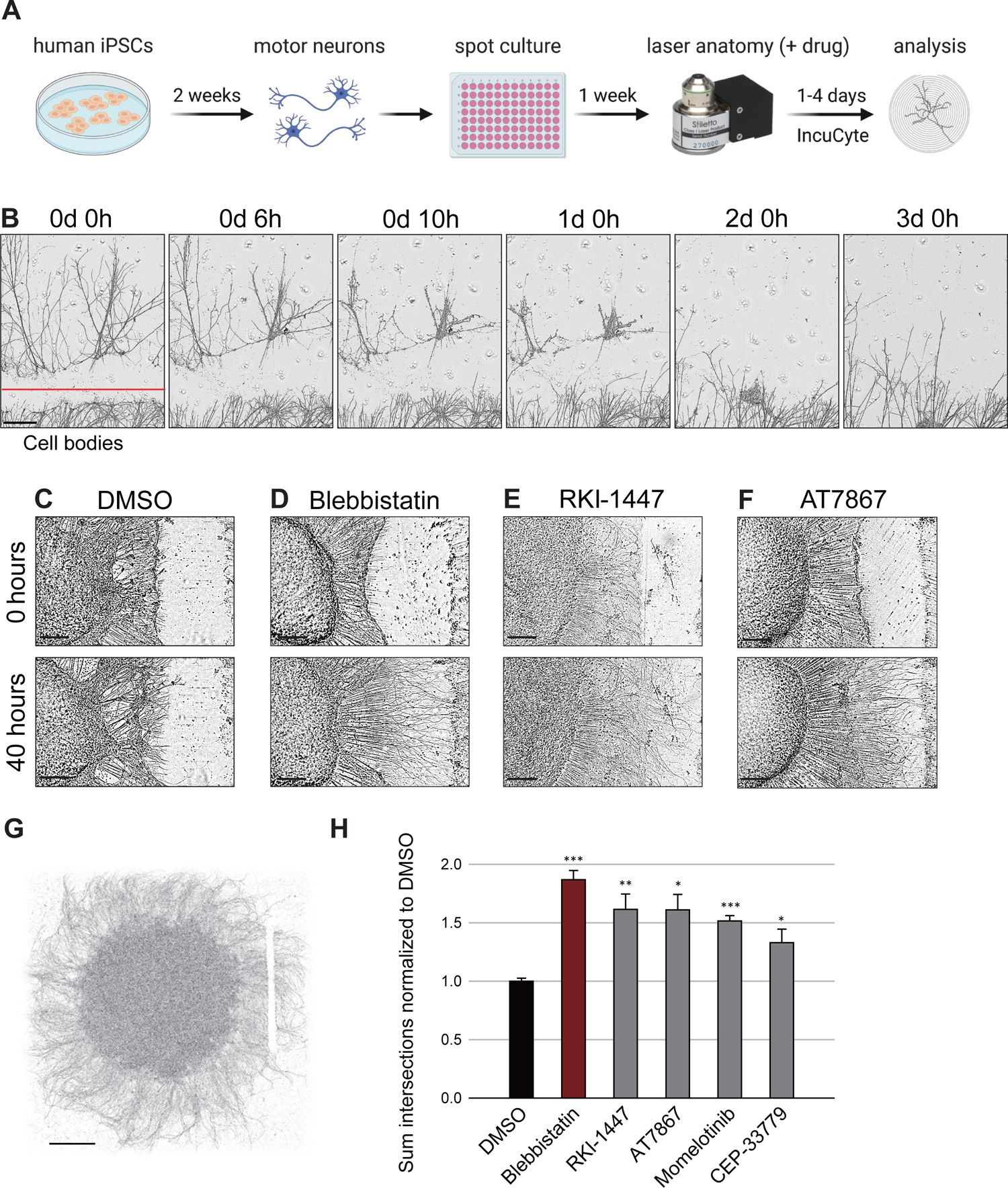
Axon regeneration in an injured human neuron spot culture model. (A) Flow chart of the human neuron spot injury assay. Human iPSC derived motor neurons were plated at high density as a spot (150,000 cells/spot). After growing for 1 week, axons were cut with laser at a specified area, and regrowth quantified with an IncuCyte. (B) Example of the response of human motor neurons to laser axotomy, red line indicates laser cutting site. Cell bodies were located below the images. Distal axons started to degenerate a few hours after injury and proximal axons regrew upward over time. Scale bar, 200μm. (C-F) Representative images of laser axotomized spots treated with DMSO (C), blebbistatin (D), RKI-1447 (E), and AT7867 (F) at time 0 and 40 hours after axotomy. Scale bar, 200μm. (G) Example of laser injured spot. Scale bar, 1mm. (H) Quantification of degree of regrowth using sholl analysis, showing regeneration response after injury and effects of treatment with top 5 hits from the primary screen. Data represented as mean ± SEM. (P=1.45E-13, 0.0084, 0.016, 0.00003, 0.0246 for blebbistatin, RKI-1447, AT7867, Momelotinib, CEP- 33779, respectively, unpaired two-tailed t-test).

The laser objective, together with an automated stage and the controller software, allowed us to target the axons and cut them precisely (Supplementary movie 3). This, together with IncuCyte imaging, enabled us to study live neuronal responses to an area of defined axonal injury in real time. Several hours after the laser axotomy distal axons start to degenerate and proximal axons begin to regrow (Figure 4B).

To study whether hits from the primary replating screen, which incorporates both neurite formation and axon elongation, also promote regeneration after axotomy, we applied hit compounds to the spot cultured neurons immediately after the laser cutting and observed axonal growth responses over time (Figure 4A). Blebbistatin significantly promoted regeneration after axonal injury in the human neuron spots (Supplementary movie 4 and 5). Figure 4C and D show the axon regrowth 40 hours after injury and drug treatment (DMSO in Figure 4C; blebbistatin in Figure 4D). Representative images show axon regrowth 40 hours after injury and drug treatment (Figure 4C-F). We quantified the degree of axon regrowth by a sholl analysis. Figure 4H shows the injury responses after treatment of the top 5 hits from the primary screen. In addition to blebbistatin, the other 4 tested hits also significantly promoted regeneration of human motor neurons after axonal injury (Figure 4H) indicating that the primary screen did detect axon regeneration promoting hits.

The spot culture injury assay is not only suitable as a secondary validation assay but is also amenable for screening. We adapted the assay so that human motor neurons are plated as one spot in each well of a 96-well plate (Figure 4A; examples in Supplementary movie 6 and 7).

### Target Validation

Knock down of non-muscle myosin II, the target of blebbistatin, with *NMIIA* shRNAs promoted axon regrowth after injury in the human motor neuron spot culture (Supplementary Figure 2A and B), confirming that inhibition of the annotated target of blebbistatin is responsible for its action. A role for non-muscle myosin inhibition on regeneration was also detected in *Caenorhabditis elegans* (Supplementary Figure 2C) where knocking down nmy-1 significantly enhanced axon regeneration in *C. elegans* after laser cutting axons *in vivo* indicating that this is a general cross-species effect.

### Genes changed after blebbistatin treatment are linked with regeneration-associated genes

Blebbistatin, as a non-muscle myosin II inhibitor reorganizes actin and microtubules in growth cones, resulting in rapid axon extension (Dupraz et al., 2019; Hur et al., 2011). Although there are many actin and microtubule regulators in the compound libraries we screened, none had such a dramatic effect in promoting neurite outgrowth as blebbistatin. We therefore hypothesized that blebbistatin may play an additional role beyond regulating actin cytoskeleton at the growth cone. Inhibition of non-muscle myosin II by blebbistatin changes gene expression in human pluripotent stem cells (Walker et al., 2010), possibly through changes in actin-myosin based contractile forces which convert to changes in signaling. To test if a blebbistatin mediated alteration in gene expression may be involved in promoting regeneration in our assays, we collected dissociated human motor neurons at different time points after re-plating on CSPG and treating with blebbistatin or a DMSO control (16, 26 and 48 hours after drug treatment). Supplementary Table 1 shows the top 20 differentially expressed genes significantly upregulated after blebbistatin treatment for 16 hours (FDR<0.1). The most up-regulated differentially expressed gene is MMP-7, matrix metallopeptidase 7, with >16-fold increase in expression after blebbistatin treatment when compared with DMSO (logFC = log2 Fold Change). Some known regeneration-associated genes (RAG) are also upregulated after blebbistatin treatment, including IFITM2, one of the top 10 upregulated differentially expressed genes (Supplementary Table 1). BMP6 expression also increased after blebbistatin treatment. BMP signaling can activate a pro-regenerative transcriptional program (Zhong and Zou, 2014).

A gene ontology analysis of those genes significantly up-regulated in blebbistatin-treated human motor neurons reveals several enriched biological processes (Figure 5A-C) and molecular functions (Supplementary Figure 3A-C), including extracellular matrix disassembly/organization, chemokine activity and microtubule motor activity. We compared the differentially expressed genes after blebbistatin treatment with the well-characterized regeneration-associated gene (RAG) set involved in axon regeneration after injury (the magenta module in Chandran et al., 2016). Blebbistatin responsive genes significantly overlap with RAGs (Hypergeometric test, P-value=1.12 x 10^-8^, 5.38 x 10^-11^ and 3.16 x 10^-11^ for 16, 26, 48 hour treatments, respectively) (Figure 5D-F). Some up-regulated RAGs induced in motor neurons after blebbistatin treatment are shown as a heatmap in Figure 5G. Gene expression changes after blebbistatin treatment are linked, therefore, with RAGs, and these may be involved, together with cytoskeletal changes in the growth cone, in increasing axon regrowth after blebbistatin treatment.

**Figure 5.**
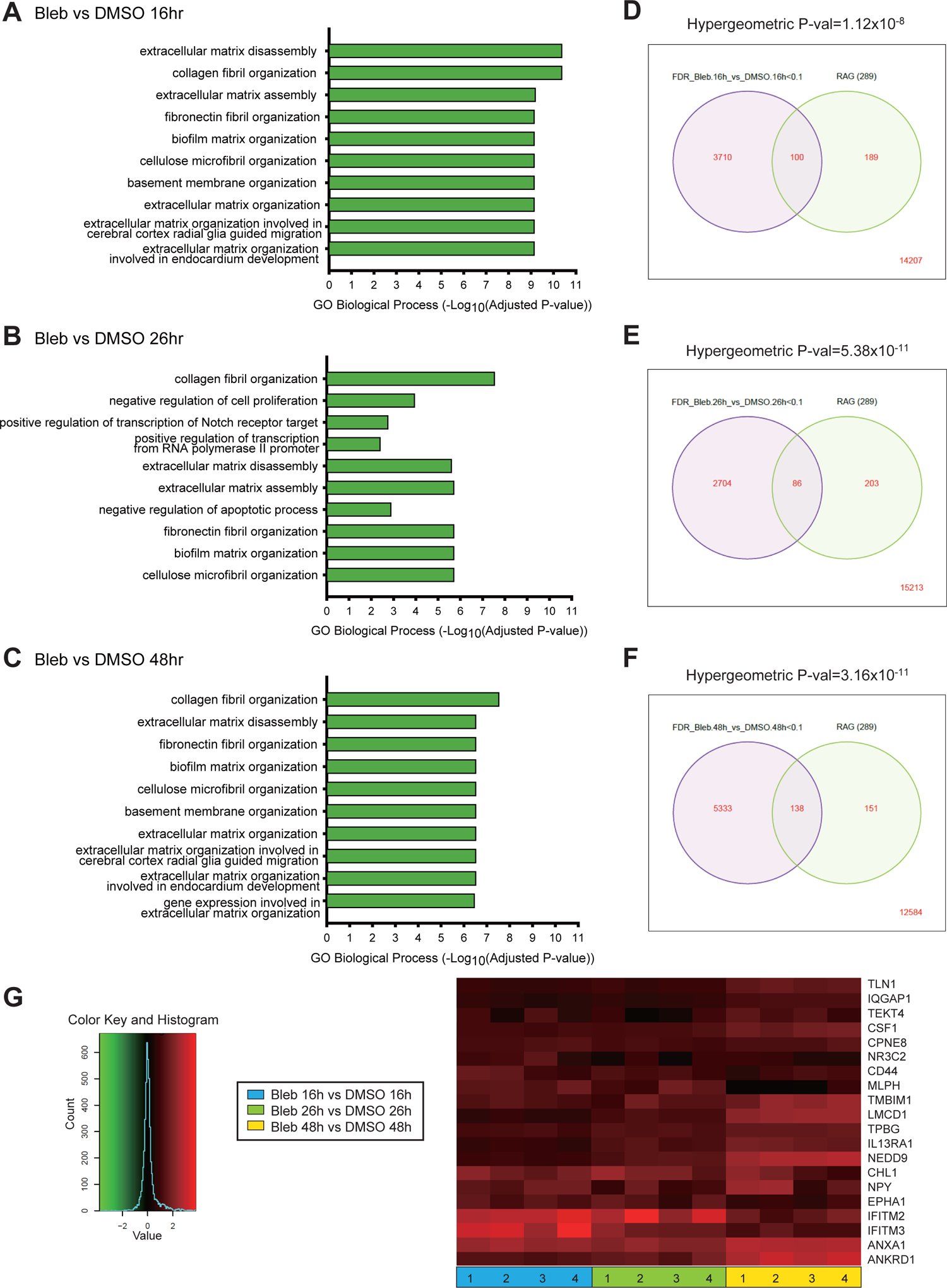
Effects of blebbistatin on transcriptional profiles of human motor neurons. (A-C) Gene Ontology enrichment scores of overlapping up-regulated genes 16 (A), 26 (B), and 48 hours (C) after blebbistatin or DMSO treatment. (D-F) Venn diagrams showing overlapping genes between blebbistatin vs DMSO and a previously defined set of regeneration associated genes (Magenta) (Chandran et al., 2016) at 16 (D), 26 (E) and 48 hours (F) after treatment. (G) A heatmap of some up-regulated regeneration associated genes after blebbistatin treatment. See also Figure S3.

### Blebbistatin promotes axon regeneration and functional recovery after sciatic nerve injury *in vivo*

To determine whether blebbistatin promotes axon regeneration after injury *in vivo*, we used a mouse sciatic nerve transection model. After transecting the sciatic nerve, the two nerve ends were bridged with a 5mm silicon tube (Figure 6A) and the extent and timing of regrowth across the bridge measured. Between the two nerve endings we left a 3mm gap and filled this with hydrogel mixed with either blebbistatin or vehicle, to locally treat the injured axons (Figure 6A and B), given that blebbistatin has poor solubility, inhibits muscle myosin II as well as non-muscle myosin II, is unstable and not well suited for systemic exposure (Dou et al., 2007; Eddinger et al., 2007; Limouze et al., 2004; Roman et al., 2018; Straight et al., 2003; Wang et al., 2008; Young et al., 2016). The nerves after regrowth were dissected at different time points, cryosectioned and immunostained with NF-200 antibody to label regenerated axons. Two weeks after injury and drug treatment, the blebbistatin treated group had more axons that had regrown through the gap region than the vehicle treated group (Figure 6C and D). Eight weeks after injury and drug treatment, the axons in the vehicle control group also grew through the gap but to a lesser extent (Figure 6F and G). A sholl analysis of axon numbers confirmed that blebbistatin significantly promoted axon regeneration (Figure 6E).

**Figure 6.**
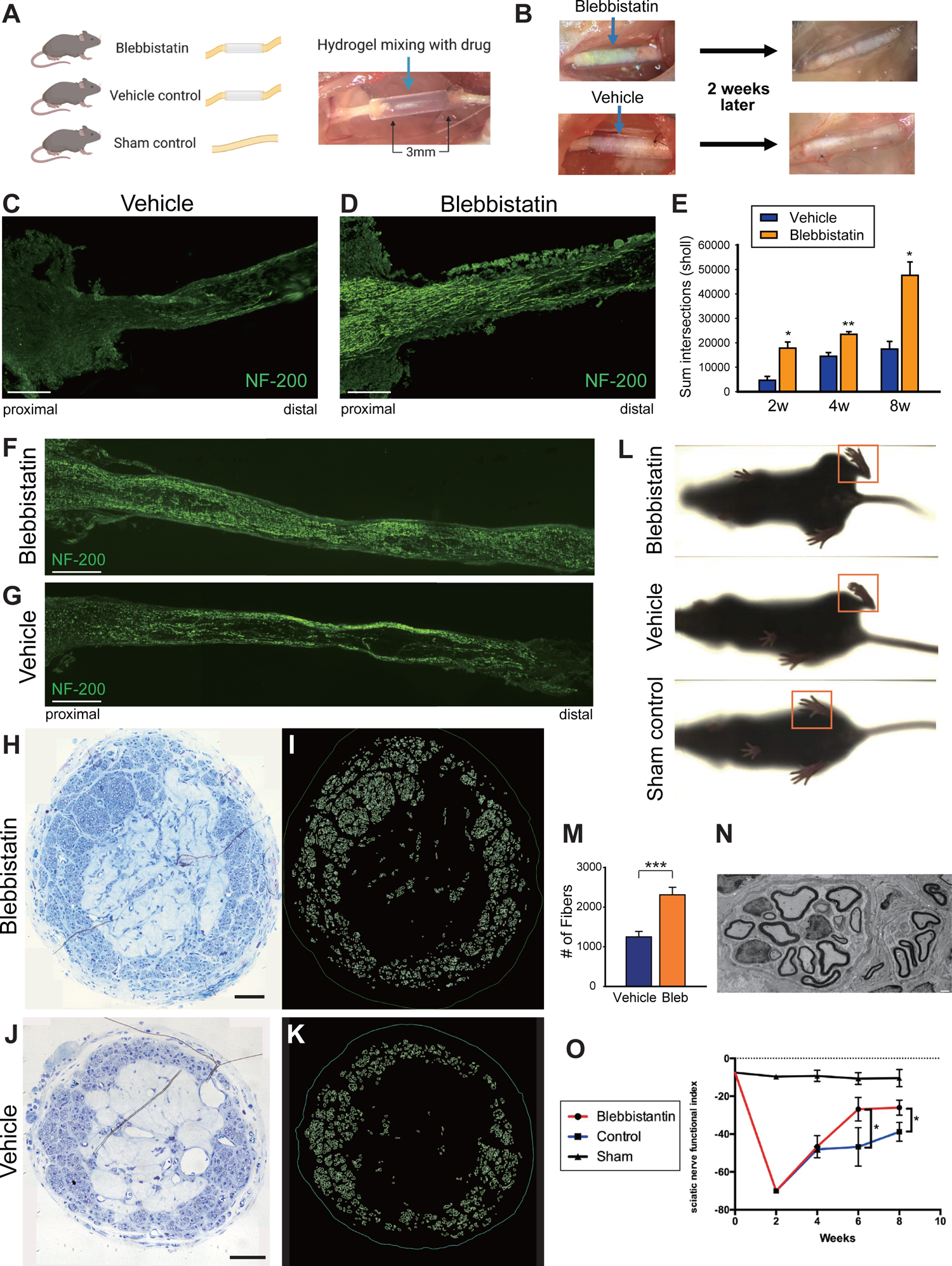
Blebbistatin promotes axon regeneration and functional recovery in an *in vivo* sciatic nerve injury model. (A) Sciatic nerve transection and bridge repair model. (B) Drug treatment (hydrogel + blebbistain or hydrogel + vehicle) in 3mm gap between nerve endings and results 2 weeks after injury. (C-G) Sciatic nerve sections 2 (C-D) or 8 weeks (F-G) after injury, stained with Neurofilament-200 antibody to label regenerating axons. Scale bar, 200μm. (E) Quantification of degree of axon regeneration by sholl analysis. Data represented as mean ± SEM. (**P-value = 0.009 for 4 weeks, *P-value = 0.0158 for 2 weeks and 0.0152 for 8 weeks, unpaired two-tailed *t*-test). (H-K) Images of representative toluidine blue-stained cross sections of nerve and segmented axons for both blebbistatin and vehicle groups. Scale bar: H, J: 50μm. (M) Note larger number of fibers in the blebbistatin group than the vehicle group (***p= 0.0007). (N) Representative electron micrograph from Blebbistatin group showing clusters of myelinated fibers. Note varied size and degree of myelination between individual fibers. (L) Digigait images of mice 8 weeks after injury and drug treatment. Red squares label paws on injured side. (O) Quantification of motor function recovery by sciatic nerve functional index of 3 groups (blebbistatin-treated, vehicle-treated, and sham) 2, 4, 6, 8 weeks after injury. Data represented as mean ± SD. (*P=0.034 for 6 weeks and *P=0.015 for 8 weeks, unpaired two-tailed t-test).

To study whether the regenerated axons remyelinated we performed light microscopy (LM) and electron microscopy (EM) on transverse sections of the regrown axons in the tube 8 weeks after the injury and blebbistatin treatment. The regenerated axons were myelinated 8 weeks after the injury both when treated with blebbistatin (Figure 6H & I) and with vehicle control (Figure 6J & K) but the blebbistatin-treated group had significantly more myelinated axons then the vehicle-treated group (Figure 6M) (P-value=0.00069, unpaired two-tailed *t*-test). The newly regenerated myelinated axons tended to form multiple distinct bundles, as shown in by EM (Figure 6N).

We also studied functional motor recovery of the blebbistatin treated mice using a DigiGait (Mouse Specifics, Inc.) to image recovery of paw spread of the mice while walking on a treadmill (Fig. 6L). After sciatic nerve transection, the hind paw on the injured side does not spread out normally (Fig. 6L, red square in the vehicle group), while the hind paw of the sham control non-injured group shows normal paw spreading (Fig. 6L, red square in the sham control group). Blebbistatin treatment improved paw spreading (Fig. 6L, red square in the blebbistatin group). Quantification of the sciatic nerve function index showed that blebbistatin improved motor functional recovery starting 6 weeks after injury (Fig. 6O).

Taken together, these data show that blebbistatin, the top hit in the human motor neuron pro-regenerative primary and secondary screens, promotes axon regeneration and functional motor recovery after sciatic nerve injury *in vivo* and that these screens can be used to find accelerators of peripheral axonal regrowth.

## Discussion

Our goal here was to develop *in vitro* human cell-based assays suitable for efficient screening in an unbiased and high throughput fashion for novel ways of promoting axon regeneration with therapeutic potential.

### Implications of the primary screening strategy

The human iPSC-derived motor neuron growth assay we developed for the primary screen comprises measuring neurite growth over a fixed period (24 hours) after re-plating differentiated motor neurons onto a CSPG substrate. The re-plating of the neurons effectively severs all their axons from the cell body and the growth that is then seen includes both initial neurite formation and then their elongation on the inhibitory substrate and is both consistent (R^2^=0.96 between replicates) and reliable (Z’ factor >0.5). While use of CSPG substrates mimics aspects of the inhibitory CNS environment (Yiu and He, 2006), it also provides a larger window to observe pro-regeneration drug effects than if the cells are plated on a permissive substrate, like laminin, as tested in pilot studies, improving the Z’.

The libraries we used for the screening comprise target-annotated bioactive compound sets, allowing us not only to identify compound hits but also targets, such as non-muscle myosin II, JAK and Akt1/2/3 and p70S6K/PKA. Involvement of non-muscle myosin II as an outgrowth suppressor target was validated by target knockdown which phenocopied the inhibitor.

Since we performed the screen in human motor neurons, one question is whether the pro-regenerative targets we identified may be applied to other neurons. To address this, we tested the effect of blebbistatin on a different neuronal type. Consistent with the effect on human motor neurons, blebbistatin also promotes outgrowth of human iPSC-derived sensory neurons (Supplementary Figure 1C), indicating the effect is not specific to motor neurons.

### Blebbistatin treatment changes gene expression in human motor neurons

Since blebbistatin caused such dramatic effects both on neurite outgrowth and axon regrowth (Figure. 2-4), we wondered if in addition to regulating the cytoskeleton at the growth cone, blebbistatin also changed gene expression, turning on genes involved in axon regeneration? To study blebbistatin induced transcriptional changes we collected human induced motor neurons after treating with blebbistatin at various time points and performed RNA Seq analysis. The top up-regulated differentially expressed gene after blebbistatin treatment for 16 hours, was matrix metallopeptidase 7 (MMP-7), with >16 fold increase in expression. MMP-7 is a secreted protease that cleaves extracellular matrix (ECM) components including CSPG (Wang et al., 2018). MMP-7 transcripts are upregulated 24 hours after a spinal cord injury in mice (Wells et al., 2003) and MMP-7 is proposed to have a neuroprotective effect through cleavage of pro-neurotrophins (Le and Friedman, 2012; Lee et al., 2001).

In addition to ECM components, and genes involved in organizing ECM molecules, genes that regulate inflammation and immunity were also upregulated (Supplementary Table 1). CXCL5 is a cytokine that has neurotrophic effects and promotes neurite growth of rat pelvic ganglia (Zhang et al., 2011). CXCL5 binds to the chemokine receptor CXCR2, which is expressed by human iPSC-induced motor neurons as well as mouse DRG neurons (http://www.painseq.com) and the signaling induced by CXCL5 might be involved in axon regrowth.

### Blebbistatin promotes axon regeneration *in vivo*

Blebbistatin significantly promoted regeneration after axonal injury in the human motor neuron spot axotomy model (Figure. 4D and H). Furthermore, blebbistatin also promoted regeneration after nerve injury *in vivo* in the adult mouse (Figure. 6D, E, F). In the sciatic nerve transection and bridge repair model, we can treat the injured nerve locally by mixing the drug with hydrogel and applying it into to the gap in the bridge between the proximal and distal parts of the nerve (Figure. 6A). Remyelination and formation of nodes of Ranvier were observed on regenerated axons crossing the gap indicating that not only was axon growth promoted but interactions with Schwann cells occur (Figure. 6H-M and data not shown). Furthermore, blebbistatin not only enhanced axon regrowth but also promoted a measurable increase in functional motor recovery after sciatic nerve injury (Figure. 6O).

While blebbistatin produces dramatic effects on axon regrowth, it is not an ideal drug for therapeutic use. It also inhibits cardiac-, skeletal-, and smooth-muscle myosin II (Dou et al., 2007; Eddinger et al., 2007; Limouze et al., 2004; Straight et al., 2003; Wang et al., 2008), has poor water solubility and poor microsomal stability (Roman et al., 2018; Young et al., 2016). Nevertheless, mice given systemic blebbistatin by intraperitoneal injection show normal behavior including locomotion and other general health measures (Young et al., 2016). could be suitable to explore promotion of CNS regeneration. Although modifications have been made to improve certain properties of blebbistatin, including solubility and photochemical features (Kepiro et al., 2014; Roman et al., 2018; Varkuti et al., 2016), a MedChem project to generate more specific non-muscle myosin II inhibitors with high potency and stability and desirable drug-like properties are required to fully explore the pro-regenerative therapeutic opportunities of this target. Utilization of the human axon outgrowth models described here, would assist such a MedChem drug development program.

### Limitations of study

There are some limitations to our primary screen, for example, we used a fixed concentration of the 4557 compounds (10μM, except for the Biomol library where concentrations vary, based upon potency and availability). Therefore, we may miss candidates whose working doses are at a different range. To address this, compound libraries could be screened at multiple concentrations, although there is always a tension between more compounds or more concentrations. We chose to conduct a full dose response analyses only on the hits.

Another limitation of the screen was that the iPSC derived motor neurons are relatively immature. Therefore, we validated the top hit in adult mice to establish if the effects were pro-regenerative rather than developmental growth-promoting, but it would be ideal to be able do this *in vitro*. In addition to the short-term neurite outgrowth promotion detected by the primary screen, we performed secondary assays that were designed to study dynamic drug effects over days and whether responses after axonal injury are similar to that after cell dissociation. Because each assay detects different aspects of the growth response and constitute independent replication of axon growth promotion actions, we feel that replication in both is informative of action on axon growth.

In conclusion, phenotypic screens of human motor neurons for axon outgrowth and regrowth, can be used for the development of treatments for the repair of the injured peripheral nervous system.

## Acknowledgments

This work was funded by the NIH (R35NS105076, C.J.W.), the Dr. Miriam and Sheldon G Adelson Medical Research Foundation (L.A.H., D.H.G. and C.J.W.), a Leonard and Isabelle Goldenson fellowship and an Ellen R. and Melvin J. Gordon fellowship (T.S-Y.H.). Figures 1J, 4A, and 6A were made using BioRender. We are grateful to Zigang He for discussions about this project and to our colleagues Andrew Snavely, Bhagat Singh, Yung-Chih Cheng, Laurel Heckman and Xuan Huang who provided expertise and advice that greatly assisted the research. We express our gratitude to Jennifer Smith and the ICCB-Longwood Screening Facility at Harvard Medical School.

## Author contributions

Conceptualization, T.S-Y.H. and C.J.W.; methodology, T.S-Y.H., T.H., Z.X.; validation, T.S-Y.H., T.H., Z.X., N.P.B., D.G.T., and X.C.; formal analysis, T.S-H.A.J.; resources, L.B.B., C.V.G., L.A.H., D.H.G. and C.J.W.; original draft, T.S-Y.H. and C.J.W.; supervision, T.S-Y.H., L.A.H., D.H.G. and C.J.W.

## Materials and methods

### Human iPSC culture

iPSCs were maintained in StemFlex medium (Thermo Fisher) on culture dishes coated with Matrigel (Corning). Cells were passaged when at approximately 80% confluence, using ReLeSR (Stem Cell Technologies) enzyme free passaging reagent. After 10 passages, a new vial of iPSCs was thawed. Human iPSC line SAH-0047 was provided by the Sahin laboratory (Boston Children’s Hospital). Human iPSC line 11a was provided by the Eggan lab (Harvard University) (Boulting et al., 2011). SAH-004 was used for the primary screen. Both SAH-004 and 11a lines were used for secondary assays and showed consistent and reproducible results.

### Motor neuron differentiation from human iPSC

iPSCs were differentiated into motor neurons as described (Klim et al., 2019) and illustrated in Figure 1J. In brief, iPSCs were dissociated into single cells using Accutase (Stem Cell Technologies) and plated at a density of 40,000 cells per cm^2^ in a 10cm Matrigel (Corning) coated dish in StemFlex (Thermo Fisher) media with 10μM Y-27632 (Tocris) on day −2. The next day, the medium was replaced by StemFlex medium without Y-27632. From day 0-5 (differentiation phase 1), the medium was changed to differentiation medium (½ Neurobasal-A (Thermo Fisher) ½ DMEM-F12 (Thermo Fisher) supplemented with ×1 B-27 supplement (Thermo Fisher), ×1 N-2 supplement (Thermo Fisher), ×1 Gibco GlutaMAX (Thermo Fisher) and ×1 non-essential amino-acids (Thermo Fisher)) supplemented with 1 µM retinoic acid (Sigma-Aldrich), 1 µM SAG (Cayman Chemical), 100 nM LDN-163189 (Sigma-Aldrich) and 10 µM SB431542 (Cayman Chemical). In phase 1, neural induction occurred with dual SMAD inhibition (SB431542 and LDN). Retinoic acid and SAG patterned the motor neuron fate. From day 6 – 13 (differentiation phase 2), the same differentiation medium was added, however, supplemented with 1 µM retinoic acid, 1 µM SAG, 4 µM SU-5402 (Sigma-Aldrich) and 5 µM DAPT (Tocris). SU-5402 and DAPT specified motor neuron identity.

On day 14, iPSC-derived motor neurons were dissociated with Accutase for 45 minutes at 37 °C, sorted by magnetic-activated cell sorting (MACS) with antibody against human CD56 neural cell adhesion molecule 1 (BD Biosciences) and anti-R-phycoerythrin (PE) magnetic particles - DM (BD Biosciences), and then re-plated on desired plates for various studies. For primary screen, 8000 neurons were seeded in each well of CSPG (0.2 ng/ml, Millipore Sigma)-coated Corning BioCoat 96-well plates (Thermo Fisher). Human induced motor neurons were then cultured in motor neuron culture media (Neurobasal-A with N2 (1:100)/ B27 (1:50)/ Glutamax (1:100)/ ×1 NEAA (1:100)/penicillin-streptomycin (1:100)) supplemented with growth factors (35μg/mL ascorbic acid (Sigma), 10ng/mL recombinant human BDNF (Thermo Fisher), 10ng/mL recombinant human GDNF (Thermo Fisher), 10ng/mL recombinant human CNTF (Thermo Fisher)).

### Compound library

The compound libraries used for the primary screen were provided by ICCB-Longwood Screening Facility (Harvard Medical School). The small molecule libraries comprised known bioactives, including Biomol ICCBL Known Bioactives 2012 Library, Mechanism of Action Library, Prestwick 3 collection, and Selleck bioactive library. The drug concentration used for the primary screen was 15μM except for Biomol library, where compound concentration varied up to 0.1 mM based upon potency and availability.

### Neurite outgrowth analysis - Primary Screen

Dissociated human iPSC-derived motor neurons were re-plated on poly-D-lysine (PDL; 100 µg/ml; Sigma-Aldrich) and CSPG-(0.2 ng/ml) coated 96 well plates at a density of 8000 cells/well and treated with compound libraries. After neurons were cultured and treated with compounds for 24 hours, neurons were fixed with 4 % paraformaldehyde (Electron Microscopy Sciences) and stained using a mouse antibody against βIII-tubulin (Sigma) and Hoechst 33342 (Thermo Fisher) for nuclear staining. Imaging and neurite outgrowth analysis were performed by a Cellomics ArrayScan XTI High-Content Screening System (Thermo Fisher). Images were acquired with a 10x objective with ArrayScan high-resolution camera. Data including valid neuron count, cell body area, total/individual neurite length, total neurite length per neuron, number of branches, etc. were collected and analyzed with the “Neuronal profiling” application. All results were normalized to DMSO control. Dose-response validation: At least 6 doses of each test compound were prepared with serial dilutions (ranging from nM to µM covering the working dose from the primary screen). For each compound concentration, four replicate wells were assayed. Cells were incubated in test compounds for 24 hours and then fixed with 4% paraformaldehyde. After staining with mouse anti-βIII-tubulin antibody and Hoechst, total neurite length per neuron and valid neuron number were analyzed using ArrayScan XTI.

### Human neuron spot culture

High density (150,000 cells per spot) of human iPSC-induced neurons were plated on completely dry PDL- (100 µg/ml) and laminin-(10 µg/ml; Thermo Fisher) coated 96-well or 6-well plates. For each spot, 1 µl cell suspension at a concentration of 150,000 cells/µl was added to the center of each well of 96-well plates or 6 µl cell suspension of 150,000 cells to each well of 6-well plates. The 1 µl spots were kept in a humidity chamber in a 37 °C incubator for 10 minutes (or 30 minutes for 6 µl spots) to settle down and attach to the plates. Then the motor neuron culture media with growth factors was slowly and carefully filled into the wells without disturbing the spots (100 µl per well for 96-well plates and 2ml per well for 6-well plates). Half medium was refreshed every 3 days. The axons grew outward radially from the spot (cell bodies) and extended continuously over time.

### Axon Injury with laser and regeneration analysis in human neuron spots

The spots were cultured for 7 days to allow motor neurons to extend long axons before a laser axotomy with a 300 mW Stiletto infrared laser (Hamilton Thorne) combined with a 20x objective. The laser module has a high-speed micro controller and an automated motorized stage. Axons were cut (as illustrated in Figure 4G) 400µm away from the cell bodies at the edge of the spot. The spots were imaged with an IncuCyte S3 Live-Cell Analysis System (Sartorius) every 3-5 hours to record the dynamic regrowth of axons. The degree of axon regeneration was analyzed using a Sholl analysis (Fiji Is Just ImageJ) (Schindelin et al., 2012). The sholl analysis quantified the sum of intersections of regenerated axons and concentric circles. The images taken right after cut (0 hour) were used to determine the starting point for the sholl analysis. All results were normalized to a control DMSO-treated group.

### RNA-seq library preparation and sequencing

Dissociated human motor neurons treated with blebbistatin or DMSO were harvested at different time points (16 hours, 26 hours and 48 hours) after re-plating and drug treatment. Total RNAs were extracted from these samples using Buffer RLT Plus (Qiagen), and then purified using RNeasy Plus mini kit (Qiagen). RNA-sequencing was carried out by HiSeq4000. Library was prepared using TruSeq Stranded RNA (100 ng) + RiboZero Gold. Quality control (QC) was performed on base qualities and nucleotide composition of sequences, to identify problems in library preparation or sequencing. Reads were trimmed and filtered, if necessary, after the QC before input to the alignment stage. Reads were aligned to the latest human GRCh38 reference genome using the STAR (v2.4.0) spliced read aligner. Average input read counts were 66.7M per sample (range 51.7 to 85.3M) and average percentage of uniquely aligned reads was 90.5% (range 88.03% to 92.0%). Total counts of read-fragments aligned to known gene regions within the human GRCh38 refSeq reference annotation were used as the basis for quantification of gene expression. Fragment counts were derived using HTS-seq (v.0.6.0) program using GRCh38 refSeq genes as model. Low count transcripts were filtered, and count data were normalized using the method of trimmed mean of M-values (TMM). Differentially expressed genes (FDR < 0.1) were then identified using the Bioconductor package EdgeR using paired-analysis (Robinson et al., 2010). The scripts that used in the RNA sequencing analyses are available at https://github.com/icnn/RNAseq-PIPELINE.git.

### Mice

10-week-old C57BL/6J mice were obtained from Jackson Laboratory (Bar Harbor). Animals were housed and handled in accordance with protocols approved by the institutional animal care and use committee (IACUC) of Boston Children’s Hospital.

### Surgery

All surgeries were performed aseptically under 2.5% isoflurane. The sciatic nerve was exposed at mid-thigh level on the left side of the mice, and then was transected using microdissection scissors. A 3mm nerve segment was removed. The two nerve endings were sutured to a 5mm SILASTIC silicone tubing (Dow Corning) with 10-0 polypropylene sutures (Ethicon), leaving a 3mm gap in between. Lifeink 200 Hydrogel (highly concentrated type 1 collagen bioinks (Advanced BioMatrix)) mixing with blebbistatin (+/-) (160μg per mice; Enzo Life Sciences) or vehicle control (6%DMSO, 44% saline, 50% hydrogel) was filled into the gap between two nerve endings with 30-gauge needles (BD). The surgical incisions were closed with 6-0 sutures (Ethicon).

### Quantification of axon regeneration *in vivo*

After sciatic nerve transection and bridge repair, sciatic nerves were dissected 2, 4 and 8 weeks after injury. Sciatic nerves were fixed and sectioned (details below in Immunohistochemistry section), and regenerating axons were visualized by staining with a chicken anti-Neurofilament H antibody (Millipore Sigma). Regenerating axons were quantified at specified distances from the injury site using the Sholl analysis plugin of Fiji ImageJ.

### Light and electron microscopic analysis

Mice were anesthetized with an overdose of ketamine and xylazine (25 mg/ml ketamine and 2.5 mg/ml xylazine in PBS) and a transcardial perfusion was performed with phosphate-buffered saline (PBS), followed by 2% paraformaldehyde, 2.5% glutaraldehyde solution (2%PF, 2.5%Glut) diluted in phosphate buffer. Sciatic nerves were dissected out and post-fixed in 2%PF, 2.5%Glut solution overnight at 4°C. Next, the tissues were rinsed in PBS and fixed in 1% Osmium (OsO_4_) solution, washed with ddH_2_O and dehydrated in a series of ethanol and 100% propylene oxide. The nerves were then embedded in Epon plastic resin, trimmed and cross sectioned (0.5μm), and stained with 1% toluidine blue solution for light microscopy. The complete sciatic nerve for each subject was photographed at 100X magnification with a Nikon Eclipse E600 microscope coupled with a Nikon camera DS-Fi3 and images automatically stitched together with Nikon NIS-Elements software. To calculate the number of fibers for the Vehicle control and Blebbistatin groups, the full cross section for each nerve was analyzed. The number of fibers was determined based on data segmentation using Neurolucida 360 software (MBF Bioscience). Statistical analysis was performed using the non-parametric Mann-Whitney test (GraphPad Prism, version 8.4.1), and considered significant when the P value was <0.05. The data are shown as mean± SE. Additional analysis was performed at the electron microscopy level. Ultrathin sections (70- 90nm) were collected on single-hole formvar-coated grids and counterstained with uranyl acetate and lead citrate. The samples were analyzed under a Tecnai G2 Spirit Twin, FEI®, ThermoFisher Scientific® or Hitachi® H-7500 electron microscopes and images were captured with NanoSprint12 AMT® camera.

### Assessment of motor function recovery

To assess motor recovery, mice were recorded using a DigiGait apparatus (Mouse Specifics). Mice were forced to walk on a transparent treadmill belt and the digital paw prints were recorded from below using a high-speed digital video camera with the information including print length (PL), toe spread (TS), and intermediate toe spread (IT). The sciatic nerve functional index (SFI) was calculated using the formula: SFI=109.5(ETS-NTS)/NTS-38.3(EPL-NPL)/NPL+13.3(EIT-NIT)/NIT-8.8 (Inserra et al., 1998). E is experimental group, N is normal group. Experiments were conducted fully blinded to drug treatment and surgery.

### Immunohistochemistry

Sciatic nerves were harvested and fixed with 4% PFA on ice for 1 hour and then cryoprotected with 30% sucrose in PBS overnight at 4°C. Sciatic nerves sections (20 μm) were collected, blocked and permeabilized with PBS containing 0.3% Triton X-100 and 10% goat serum (PBTGS) for 1 hour at room temperature (RT). Sections were incubated with a chicken anti-Neurofilament H antibody (AB5539, Millipore Sigma; 1:1000) diluted in PBTGS overnight at RT and then incubated with Alexa Fluor 488 goat antibody against chicken IgY (A11039, Thermo Fisher; 1:1000) and Hoechst 33342 (H3570, Thermo Fisher; 1:2000) diluted in PBTGS for 1 hr at RT. Images were acquired using a Nikon Eclipse 80I Microscope.

Human induced motor neuron culture were fixed with 4% PFA for 1 hour, washed with PBS and then permeabilized with PBS containing 0.3% Triton X-100 and 10% goat serum (PBTGS) for 1 hour at RT. Neurons were then incubated with a mouse monoclonal antibody against βIII-tubulin (T8660, Sigma; 1:1000) diluted in PBTGS overnight, and then were incubated with Alexa Fluor 568 goat anti-mouse IgG (A11031, Thermo Fisher; 1:1000) and Hoechst 33342 (1:2000) diluted in PBTGS for 1 hour at RT. Images were acquired using ArrayScan XTI high-resolution camera (Thermo Fisher).

### Caenorhabditis elegans study

B. elegans strain TU3568 (sid-1(pk3321) him-5(e1490) V; lin-15B(n744) X; uIs71 [(pCFJ90) pmyo-2::mCherry + pmec-18::sid-1]) was employed for neuronal-specific RNAi knockdown of the C. elegans homologs of non-muscle myosin II, nmy-1 and nmy-2 (Calixto et al., 2010; Taub et al., 2018). Clones from the Vidal library were streaked onto LB plates containing carbenicillin and grown overnight at 37°C. Colonies were picked and grown overnight at 37°C in 6mL of LB supplemented with carbenicillin. 200µL of this subculture was then spread over a NGM agar plate containing 2mM IPTG to induce RNAi expression. A bacterial lawn was grown at room temperature for 3 days. Gravid adult hermaphrodites were bleached and embryos allowed to crawl onto the RNAi-containing bacteria (termed F1s). 30 F1s were then picked onto a new RNAi-containing plate and allowed to lay eggs for 4 hours. The resulting embryos (F2s) were examined in regeneration assays at the young adult stage. Regeneration between control RNAi and nmy RNAi were examined.

Axotomy was performed on a Nikon Ti-2000 inverted fluorescent microscope coupled with a Ti:Sapphire infrared laser (Mantis Pulse Switch, Coherent Inc.). Day 1 adults were placed on a 10% agarose pad and immobilized with a slurry of polystyrene beads (Polysciences). A 1KHz train of 100 fs pulses (15-30 nJ/pulse) was focused through a Nikon 60X 1.4 NA objective to vaporize specifically at the focal point, targeted to be at 20µm and 40µm from the cell body of the Anterior Lateral Microtubule (ALM) neuron, thus creating a gap of ∼20µm at the site of injury. This procedure was performed in a span of 15 minutes before returning the adults to the RNAi agar plates for 24 hours. Regeneration was examined by immobilizing the adults with 5mM sodium azide and capturing a Z-stack a Nikon 40X 1.3NA. A maximum projection image was made and regeneration lengths from the site of injury were measured with FIJI.

#### Statistical Analysis

Data were statistically tested with unpaired unequal variances student t-tests. Results in Figure 1, 2 and 6 are expressed as mean ± standard deviation. Data in Figure 3 and 4 are expressed as the mean ± standard error of the mean. In the figures, the significance is indicated with asterisks (*for p<0.05, ** for p<0.01 and *** for p<0.001).

## Supplemental information

**Supplementary Table 1:**
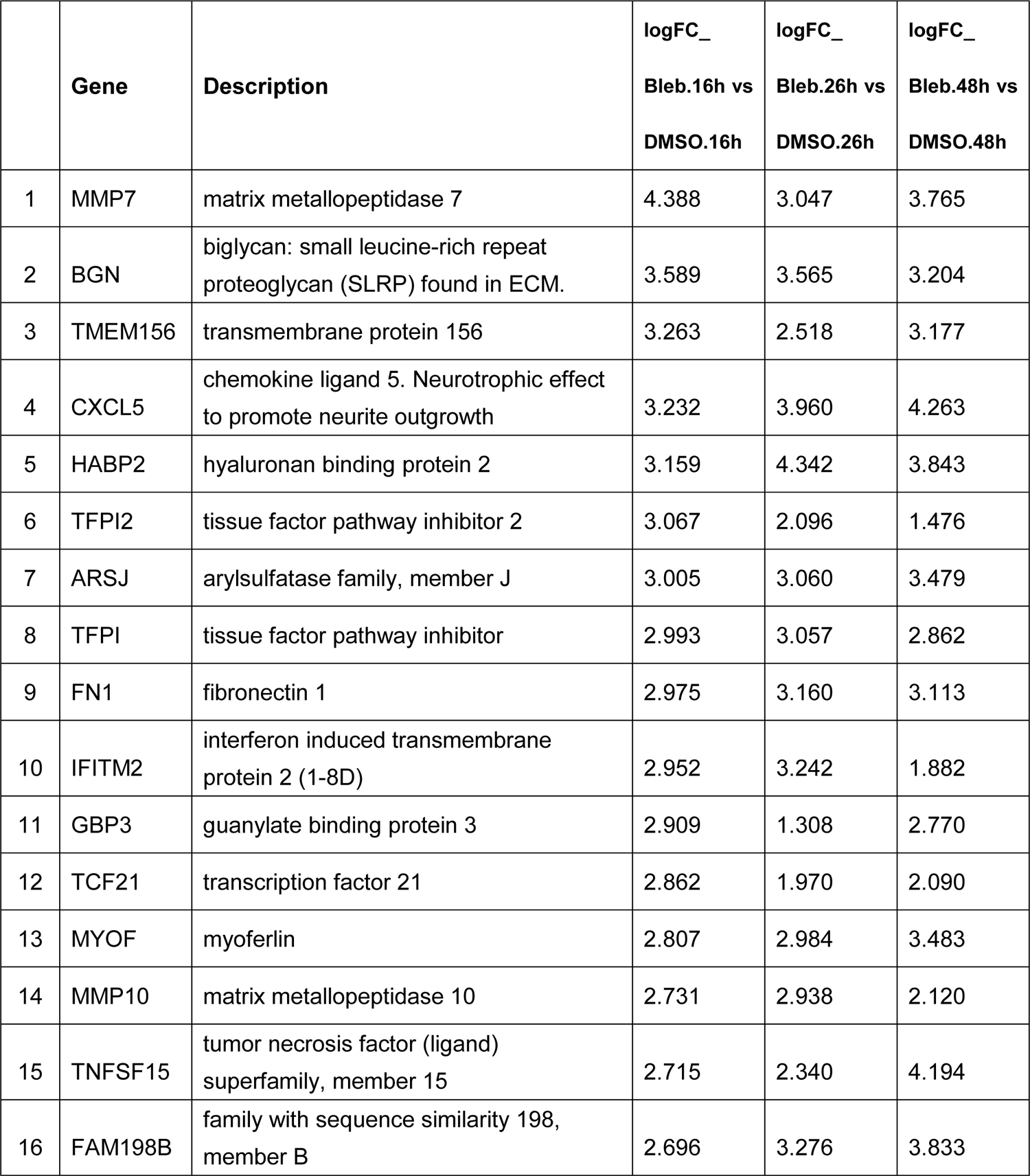
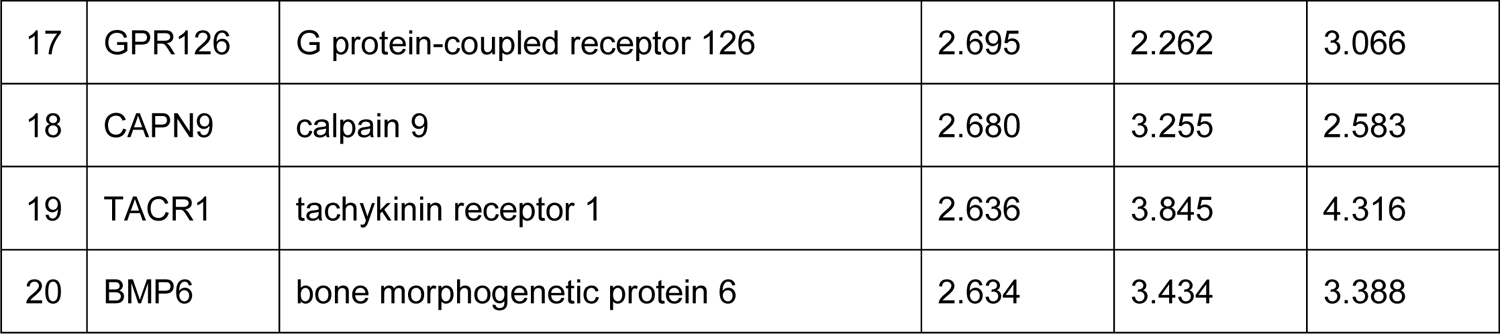
Top 20 differentially expressed genes significantly upregulated after blebbistatin treatment (FDR&0.1).

Supplementary Figure 1. Dose response curves of two JAK inhibitors. Blebbistatin promotion of outgrowth of human iPSC-derived sensory neurons. (A-B) Dose response curves of two JAK inhibitors, Baricitinib (A) and Ruxolitinib (B), showing their effects on promoting neurite outgrowth of human motor neurons after a 24 hour treatment. (C-D) The effect of blebbistatin on neurite outgrowth after re-plating on a permissive substrate, laminin. (C) Human iPSC-derived sensory neurons; (D) human iPSC-derived motor neurons. Note that human induced sensory neurons did not survive on CSPG after re-plating. Data are represented as mean ± SD. (**P=0.0027, ***P=1.0018e-10, unpaired two-tailed t-test).

Supplementary Figure 2. Target validation: knockdown of non-muscle myosin II promotes axon regeneration after injury in human neurons, and in *C. elegans*. (A-B) Knockdown of non-muscle myosin II with MYH9 shRNA (A) accelerated axon regrowth after injury in human motor neuron spot culture when compared with non-silencing control (B). (C) Knockdown of non-muscle myosin homolog nmy-1 significantly enhances regeneration in *C. elegans*.

Supplementary Figure 3. Effects of blebbistatin on transcriptional profiles of human motor neurons. (A-C) Gene ontology analysis of the genes significantly up-regulated in blebbistatin-treated human motor neurons shows enriched molecular functions 16 (A), 26 (B) and 48 hours (C) after blebbistatin vs. DMSO treatment.

