## Supplemental Information for "Screening for axon regeneration promoting compounds with human iPSC-derived motor neurons"

**Supplementary Table 1: Top 20 differentially expressed genes significantly upregulated after blebbistatin treatment (FDR<0.1)**.

| | Gene | Description | logFC_ Bleb.16h vs DMSO.16h | logFC_ Bleb.26h vs DMSO.26h | logFC_ Bleb.48h vs DMSO.48h |
|---|---|---|---|---|---|
| 1 | MMP7 | matrix metallopeptidase 7 | 4.388 | 3.047 | 3.765 |
| 2 | BGN | biglycan: small leucine-rich repeat proteoglycan (SLRP) found in ECM. | 3.589 | 3.565 | 3.204 |
| 3 | TMEM156 | transmembrane protein 156 | 3.263 | 2.518 | 3.177 |
| 4 | CXCL5 | chemokine ligand 5. Neurotrophic effect to promote neurite outgrowth | 3.232 | 3.960 | 4.263 |
| 5 | HABP2 | hyaluronan binding protein 2 | 3.159 | 4.342 | 3.843 |
| 6 | TFPI2 | tissue factor pathway inhibitor 2 | 3.067 | 2.096 | 1.476 |
| 7 | ARSJ | arylsulfatase family, member J | 3.005 | 3.060 | 3.479 |
| 8 | TFPI | tissue factor pathway inhibitor | 2.993 | 3.057 | 2.862 |
| 9 | FN1 | fibronectin 1 | 2.975 | 3.160 | 3.113 |
| 10 | IFITM2 | interferon induced transmembrane protein 2 (1-8D) | 2.952 | 3.242 | 1.882 |
| 11 | GBP3 | guanylate binding protein 3 | 2.909 | 1.308 | 2.770 |
| 12 | TCF21 | transcription factor 21 | 2.862 | 1.970 | 2.090 |
| 13 | MYOF | myoferlin | 2.807 | 2.984 | 3.483 |
| 14 | MMP10 | matrix metallopeptidase 10 | 2.731 | 2.938 | 2.120 |
| 15 | TNFSF15 | tumor necrosis factor (ligand) superfamily, member 15 | 2.715 | 2.340 | 4.194 |
| 16 | FAM198B | family with sequence similarity 198, member B | 2.696 | 3.276 | 3.833 |
| 17 | GPR126 | G protein-coupled receptor 126 | 2.695 | 2.262 | 3.066 |
| 18 | CAPN9 | calpain 9 | 2.680 | 3.255 | 2.583 |
| 19 | TACR1 | tachykinin receptor 1 | 2.636 | 3.845 | 4.316 |
| 20 | BMP6 | bone morphogenetic protein 6 | 2.634 | 3.434 | 3.388 |

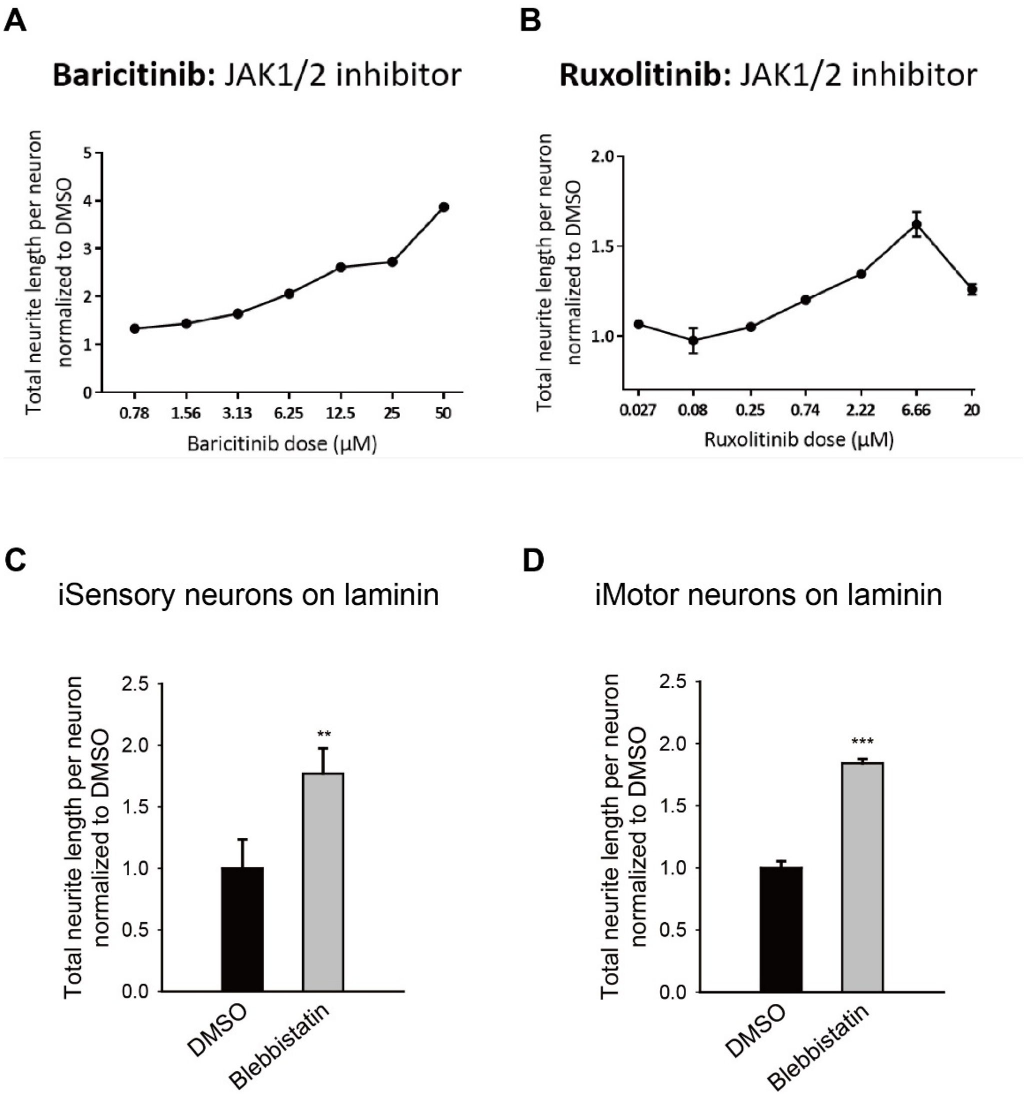


**Supplementary Figure 1. Dose response curves of two JAK inhibitors. Blebbistatin promotion of outgrowth of human iPSC-derived sensory neurons.** (A-B) Dose response curves of two JAK inhibitors, Baricitinib (A) and Ruxolitinib (B), showing their effects on promoting neurite outgrowth of human motor neurons after a 24 hour treatment. (C-D) The effect of blebbistatin on neurite outgrowth after re-plating on a permissive substrate, laminin. (C) Human iPSC-derived sensory neurons; (D) human iPSC-derived motor neurons. Note that human induced sensory neurons did not survive on CSPG after re-plating. Data are represented as mean ± SD. (**P=0.0027, ***P=1.0018e-10, unpaired two-tailed t-test).

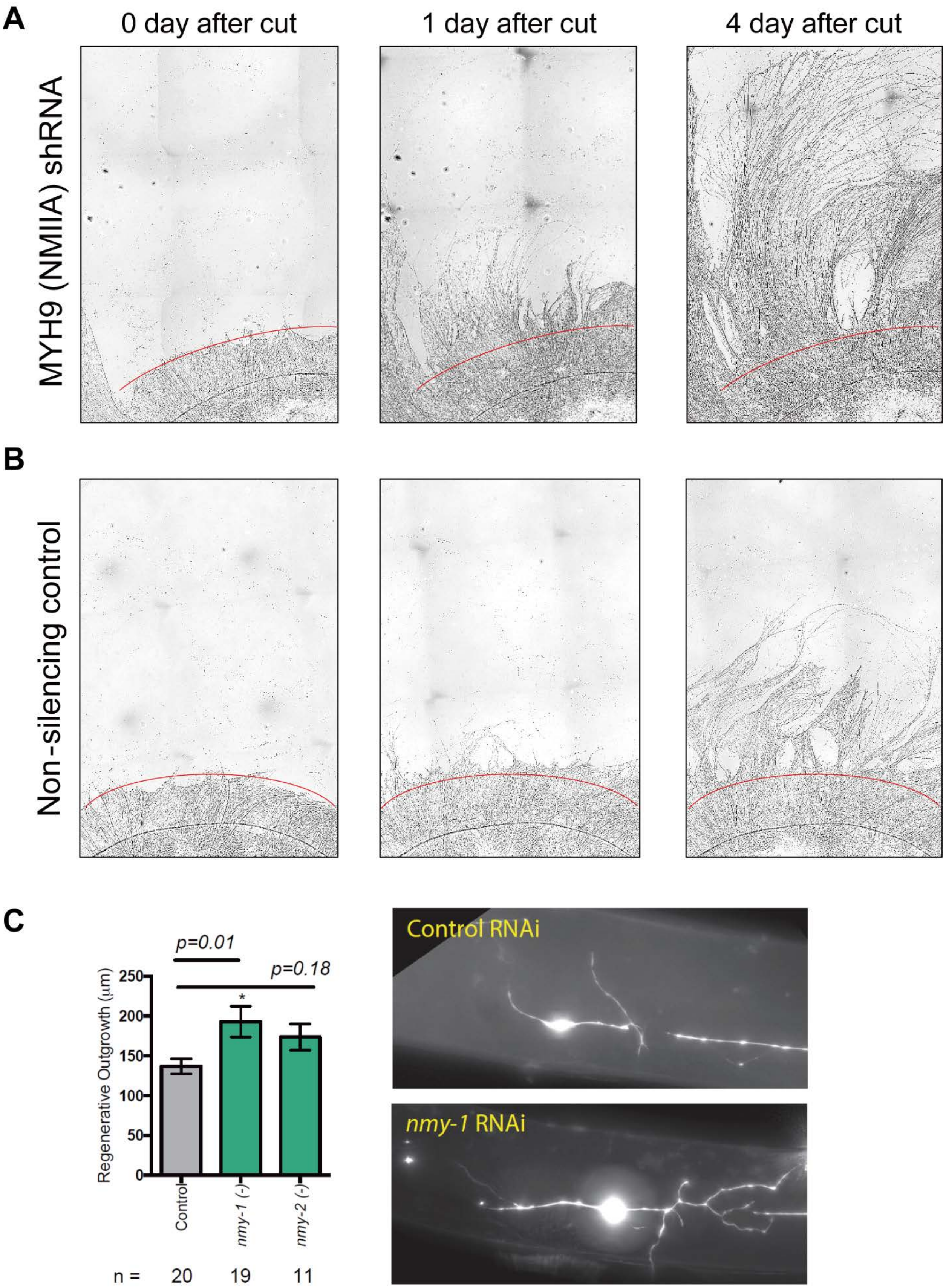
A
0 day after cut
1 day after cut
4 day after cut
MYH9 (NMIIA) shRNA
B
Non-silencing control
C
p=0.01
p=0.18
Regenerative Outgrowth (μm)
250
200
150
100
50
0
*
Control
nmy-1 (-)
nmy-2 (-)
n =
20
19
11
Control RNAi
nmy-1 RNAi

**Supplementary Figure 2. Target validation: knockdown of non-muscle myosin II promotes axon regeneration after injury in human neurons, and in *C. elegans*.**

(A-B) Knockdown of non-muscle myosin II with MYH9 shRNA (A) accelerated axon regrowth after injury in human motor neuron spot culture when compared with non-silencing control (B). (C) Knockdown of non-muscle myosin homolog nmy-1 significantly enhances regeneration in *C. elegans*.

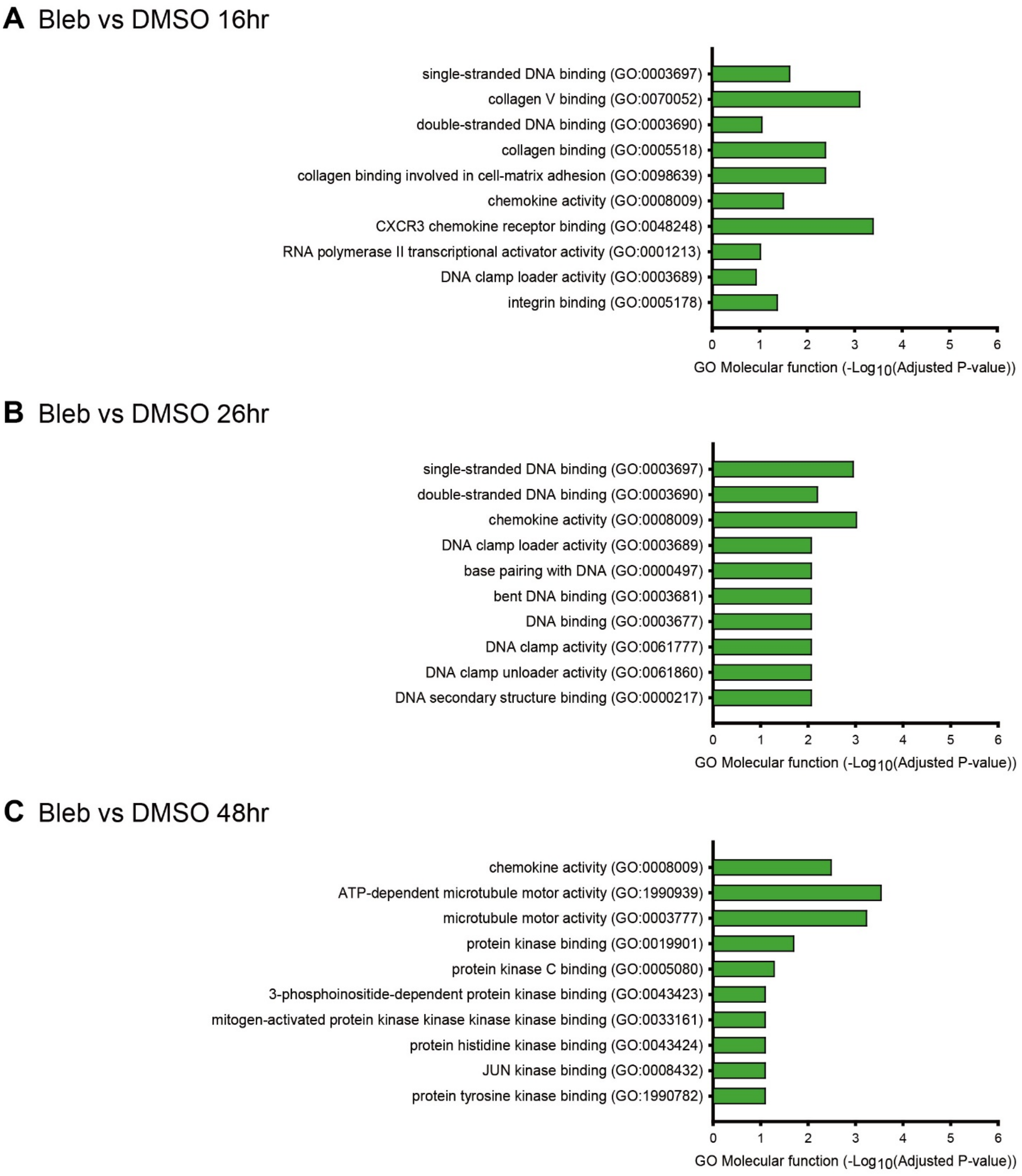


**Supplementary Figure 3. Effects of blebbistatin on transcriptional profiles of human motor neurons**.

(A-C) Gene ontology analysis of the genes significantly up-regulated in blebbistatin-treated human motor neurons shows enriched molecular functions 16 (A), 26 (B) and 48 hours (C) after blebbistatin vs. DMSO treatment.

**Movie S1. Blebbistatin treated human motor neurons over 5 days.**

**Movie S2. DMSO treated human motor neurons over 5 days.**

**Movie S3. Laser cutting axons of human induced neurons.**

**Movie S4. Response after axon injury and blebbistatin treatment in human neuron spot culture.**

**Movie S5. Response after axon injury and DMSO treatment in human neuron spot culture.**

**Movie S6. Regrowth after injury and blebbistatin treatment - whole spot view.**

**Movie S7. Regrowth after injury and DMSO treatment - whole spot view.**
